# Artificial Intelligence for Ecological and Evolutionary Synthesis

**DOI:** 10.1101/161125

**Authors:** Philippe Desjardins-Proulx, Timothée Poisot, Dominique Gravel

## Abstract

The grand ambition of theorists studying ecology and evolution is to discover the logical and mathematical rules driving the world’s biodiversity at every level from genetic diversity within species to differences between populations, communities, and ecosystems. This ambition has been difficult to realize in great part because of the complexity of biodiversity. Theoretical work has led to a complex web of theories, each having non-obvious consequences for other theories. Case in point, the recent realization that genetic diversity involves a great deal of temporal and spatial stochasticity forces theoretical population genetics to consider abiotic and biotic factors generally reserved to ecosystem ecology. This interconnectedness may require theoretical scientists to adopt new techniques adapted to reason about large sets of theories. Mathematicians have solved this problem by using formal languages based on logic to manage theorems. However, theories in ecology and evolution are not mathematical theorems, they involve uncertainty. Recent work in Artificial Intelligence in bridging logic and probability theory offers the opportunity to build rich knowledge bases that combine logic’s ability to represent complex mathematical ideas with probability theory’s ability to model uncertainty. We describe these hybrid languages and explore how they could be used to build a unified knowledge base of theories for ecology and evolution.

case study you explore using the Salix tritrophic system.

## 0 Introduction

Almost four decades ago, Ralph W. Lewis argued for the formalization of evolutionary theory and the recognition of evolution as a system of theories. In his words, “when theories are partially formalized […] the intra- and interworkings of theories become more clearly visible, and the total structure of the discipline becomes more evident” [45]. Supporting Lewis’ point, Queller recently showed how Fisher’s fundamental theorem of natural selection, Price’s theorem, the Breeder equation of quantitative genetics, and other key formulas in ecology and evolution were related [62]. In the same vein, Rice formulated an axiomatic theory of evolution based on a stochastic version of Price’s theorem [63]. These projects fall under the scope of automated theorem proving, one of the oldest and most mature branches of Artificial Intelligence [31]. Theories can be written in some formal language, such as first-order logic or type theory, and then algorithms are used to ensure the theories can be derived from a knowledge base of axioms and existing results. In the last few decades, mathematicians have built knowledge bases with millions of helper theorems to assist the discovery of new ideas [39]. For example, the Mizar Mathematical Library is a growing library of theorems, which are added after new candidate theorems are approved by the proof checker and peer-reviewed for style. Such libraries help mathematicians juggle with a growing body of knowledge and offers a concrete answer to the issue of knowledge synthesis. Mizar uses a language powerful enough for the formalization of evolutionary theories envisioned by Lewis and the result of Queller on Price’s theorem and its relationship to other theories. It is also expressive enough to build a knowledge base out of Rice’s axiomatic theory of evolution. Doing so would force us to think more clearly about the theoretical structure of evolution, with theoretical ecology facing a similar state of disorganization [45]. Case in point: theoretical community ecologists have been criticized for focusing on a single prediction for theories capable of making several [49]. An example of this is Hubbell’s neutral theory of biodiversity [36], which uses an unrealistic point-mutation model that does not fit with our knowledge of speciation, leading to odd predictions [23, 19, 20]. In logic-based (also called symbolic) systems like Mizar, all formulas involving speciation would be implicitly linked together. Storing ecological theories in such knowledge base would automatically prevent inconsistencies and highlight the consequences of theories.

Despite the importance of formalization, it remains somewhat divorced from an essential aspect of theories in ecology and evolution: their probabilistic and fuzzy nature. As a few examples: a surprisingly common idea found in ecological theories is that predators are generally larger than their prey, a key assumption of the food web model of Williams and Martinez [76]; deviations from the Hardy–Weinberg principle are not only common but tend to give important information on selective pressures; and nobody expects the Rosenzweig–MacArthur predator-prey model to be exactly right. In short, important ideas in ecology and evolution do not fit the true/false epistemological framework of systems like Mizar, and ideas do not need to be derived from axiomatic principles to be useful. We are often less concerned by whether a formula can be derived from axioms than in how it fits a particular dataset. In the 1980s, Artificial Intelligence experts developed probabilistic graphical models to handle large probabilistic systems [57]. While probabilistic graphical models are capable of answering probabilistic queries for large systems of variables, they cannot represent or reason with sophisticated mathematical formulas. Alone, neither logic nor probability theory is enough to elucidate the structure of theories in ecology and evolution.

For decades, researchers have tried to unify probability theory with rich logics to build knowledge bases both capable of the sophisticated mathematical reasoning found in automated theorem provers and the probabilistic reasoning of graphical models. Recent advances moved us closer to that goal [64, 27, 74, 54, 35, 70, 4]. Using these systems, it is possible to check if a mathematical formula can be derived from existing results and also possible to ask probabilistic queries about theories and data. The probabilistic nature of these representations is a good fit to learn complex logical and mathematical formulas from data [41]. Within this framework, there is no longer a sharp distinction between *theory* and *data*, since the knowledge base defines a probability distribution over all objects, including logical relationships and mathematical formulas.

For this article, we introduce key ideas on methods at the frontier of logic and probability, beginning with a short survey of knowledge representations based on logic and probability. First-order logic is described, along with how it can be used in a probabilistic setting with Markov logic networks [64]. We detail how theories in ecology and evolution can be represented with Markov logic networks, as well as highlighting some limitations. We present a case study involving a tritrophic system to demonstrate the strengths and weaknesses of Markov logic networks. Synthesis in ecology and evolution has been made difficult by the sheer number of theories involved and their complex relationships [59]. Practical representations to unify logic and probability are relatively new, but we argue they could be used to achieve greater synthesis by allowing the construction of large, flexible knowledge bases with a mix of mathematical and scientific knowledge.

## 1 Knowledge representations

Traditional scientific theories and models are mathematical, or logic-based. Einstein’s *e* = *mc*^2^ established a relationship between energy *e*, mass *m*, and the speed of light *c*. This mathematical knowledge can be reused: in any equation with energy, we could replace *e* with *mc*^2^. This ability of mathematical theories to establish precise relationships between concepts, which can then be used as foundations for other theories, is fundamental to how science grows and forms an interconnected corpus of knowledge. The formula is implicitly connected to other formulas involving the same symbol, such that if we were to establish a different but equivalent way to represent the speed of light *c*, it could automatically substitute *c* in *e* = *mc*^2^.

Artificial Intelligence researchers have long been interested in expert systems capable of scientific discoveries, or simply capable of storing scientific and medical knowledge in a single coherent system. *Dendral*, arguably the first expert system, could form hypotheses to help identify new molecules using its knowledge of chemistry [46]. In the 1980s, *Mycin* was used to diagnose blood infections (and did so more accurately than professionals) [14]. Both systems were based on logic, with *Mycin* adding a “confidence factor” to its rules to model uncertainty. These expert systems generally relied on a simple logic system not powerful enough to handle uncertainty. With few exceptions, the rules were hand-crafted by human experts. After the experts established the logic formulas, the systems acted as static knowledge bases and unable to discover new rules. Algorithms have been developed to learn new logic rules from data [52, 53], but the non-probabilistic nature of the resulting knowledge base makes it difficult to handle real-world uncertainty. In addition to expert systems, logic systems are used to store mathematical knowledge and perform automatic theorem proving [31]. Pure logic has rarely been used in ecology and evolution, but recent studies have shown its ability to reconstruct food webs from data [10, 72].

There are many different logics for expert systems and automatic theorem proving [31, 61, 56]. We will focus on first-order logic, the most commonly used logic in efforts to unify logic with probability. A major reason for adopting rich logics, whether first-order or higher-order, is to allow for the complex relationships found in ecology and evolution to be expressed in concise formulas. Stuart Russell noted that “the rules of chess occupy 10^0^ pages in first-order logic, 10^5^ pages in propositional logic, and 10^38^ pages in the language of finite automata” [65]. Similarly, first-order logic allows us to directly express complex ecological ideas in a simple but formal language.

In mathematics, a function *f* maps terms **X** (its domain) to other terms **Y** (its codomain) *f* : **X** → **Y**. The number of arguments of a function, |**X**|, is called its arity. The atomic element of first-order logic is the **predicate**: a function that maps 0 or more terms to a truth value: false or true. In first-order logic, terms are either variables, constants, or functions. A **variable** ranges over a domain, for example *x* could range over integers, *p* over a set of species, and *city* over a set of cities. **Constants** represent values such as 42, *Manila, π*. Lastly, **functions** map terms to other terms such as multiplication, integration, *sin, CapitalOf* (mapping a country to its capital). Variables have to be **quantified** either universally with ∀ (forall), existentially with ∃ (exists), or uniquely with ∃!. ∀*x* : *p*(*x*) means *p*(*x*) must hold true for all possible values of *x*. ∃*x* : *p*(*x*) means it must hold for at least one value of *x* while ∃!*x* : *p*(*x*) means it must hold for exactly one value of *x*. Using this formal notation, we can write the relationship between the basal metabolic rate (BMR) and body mass (*Mass*) for mammals [2]:

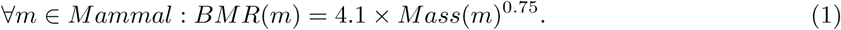

This formula has one variable *m* which is universally quantified: ∀*m* ∈ *Mammal* reads “for all *m* in the set *Mammal*”. It has two constants: the numbers 4.1 and 0.75, along with four functions (*BMR, Mass*, multiplication, exponentiation). The equal sign = is the sole predicate.

A first-order logic **formula** is either a lone predicate or a complex formula formed by linking formulas using the unary connective ¬ (negation) or binary connectives (*and* ∧, *or* ∨, *implication* ⇒, see table 1). For example, *PreyOn*(*s*_*x*_, *s*_*y*_) is a predicate that maps two species to a truth value, in this case whether the first species preys on the second species, and *IsParasite*(*s*) is a predicate that is true if species *s* is a parasite. We could also have a function *Mass*(*s*_*x*_) mapping a species to its body mass. We can build more complex formulas from there, for example:

**Table 1:**
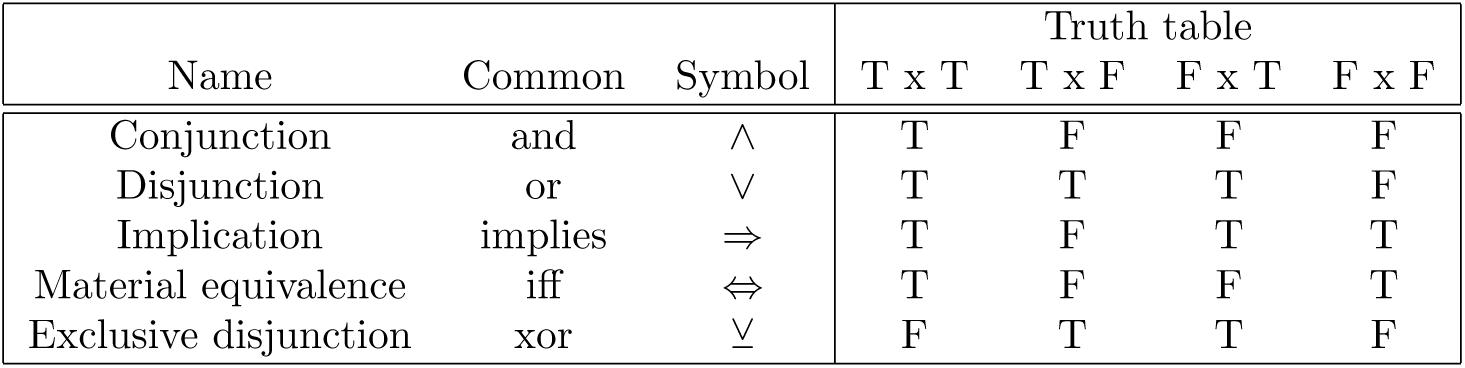
Common binary connectives. The table shows the resulting truth value (T: True, F: False) for all possible combinations. *iff* is read *if and only if*. Implication is one of the most common connective and may have surprising behavior. In particular, it will always return true when the left-side is false. While this may seem odd, it allows us to make statements such as 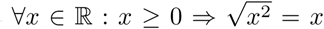. This formula holds for all real numbers, including negative ones, since with *x* = −1, *x* ≥ 0 is false and *F* ⇒ *F* returns true.

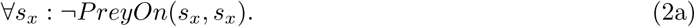

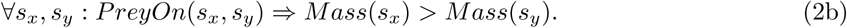

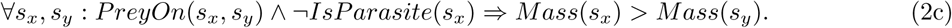

The first formula says that species don’t prey on themselves. The second formula says that predators are larger than their prey (> is a shorthand for the *greater than* predicate). The third formula refines the second one by adding that predators are larger than their prey unless the predator is a parasite. None of these rules are expected to be true all the time, which is where mixing probability with logic will come in handy. The Rosenzweig-MacArthur equation can also easily be expressed with first-order logic:

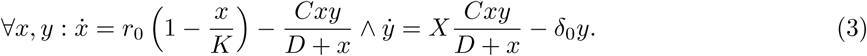

This formula has four functions: the time differential 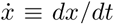, multiplication, addition, and subtraction. Prey *x* and predator *y* are universally quantified variables while *r*_0_, *K, C, D, X, δ*_0_ are constants. The formula has only one predicate, =, and both sides of the formula are connected by ∧, the symbol for conjunction (“and”).

A **knowledge base** 𝒦 in first-order logic is a set of formulas 𝒦 = {*f*_0_, *f*_1_, …, *f*_|𝒦|−1_}. First-order logic is expressive enough to represent and manipulate complex logic and mathematical ideas. It can be used for simple ideas such that predators are generally larger than their prey (eq. 2b), mathematical formulas for predator-prey systems equation (eq. 3), and also to establish the logical relationship between various predicates. We may want a *PreyOn* predicate to tell us whether *s*_*x*_ preys on *s*_*y*_, but also a narrower *PreyOnAt*(*s*_*x*_, *s*_*y*_, *l*) predicate to model whether *s*_*x*_ preys on *s*_*y*_ at a specific location *l*. In this case, it would be a good idea to have the formula ∀*s*_*x*_, *s*_*y*_, *l* ⇒ : *PreyOnAt*(*s*_*x*_, *s*_*y*_, *l*) *PreyOn*(*s*_*x*_, *s*_*y*_). Given this formula and the data point *PreyOnAt*(*Wolverine, Rabbit, Quebec*), we do not need *PreyOn*(*Wolverine, Rabbit*) to be explicitly stated, ensuring the larger metaweb [60] is always consistent with information from local food webs.

An **interpretation** defines which object, predicate, or function is represented by which symbol, e.g., it says *PreyOnAt* is a predicate with three arguments, two species and one location. The process of replacing variables with constants is called **grounding**, and we talk of ground terms / predicates / formulas when no variables are present. Together with an interpretation, a **possible world** assigns truth values to each possible ground predicate, which can then be used to assign truth values to a knowledge base’s formulas. *PreyOn*(*s*_*x*_, *s*_*y*_) can be neither true nor false until we assign constants to the variables *s*_*x*_ and *s*_*y*_. Constants are typed, so a set of constants 𝒞 may include two species {*Gulo gulo, Orcinus orca*} and three locations {*Quebec, Fukuoka, Arrakis*}. The constants 𝒞 yield 2^2^ × 3 possible ground predicates for *PreyOnAt*(*s*_*x*_, *s*_*y*_, *l*):

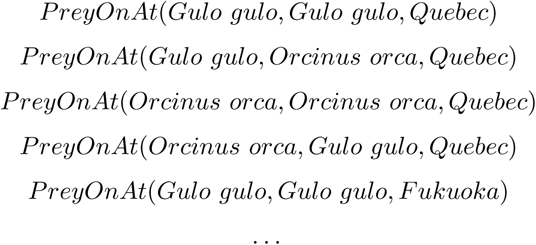

and only two possible ground predicates for *IsParasite*:

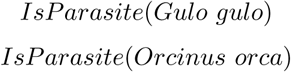

We say a possible world **satisfies** a knowledge base (or a single formula) if all the formulas are true given the ground predicates. A basic question in first-order logic is to determine whether a knowledge base 𝒦 **entails** a formula *f*, or 𝒦 |= *f*. Formally, the entailment 𝒦 |= *f* means that for all possible worlds in which all formulas in 𝒦 are true, *f* is also true. More intuitively, it can be read as the formula *following from* the knowledge base [66]. A **proof** in first-order logic can be derived using **inference rules** such as **Modus Ponens**:

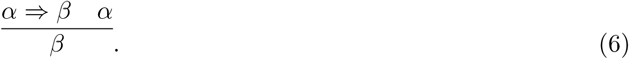

This notation reads: infer *β* if *α* ⇒ *β* is true and *α* is true. See [31] for a detailed look at inference rules in first-order logic.

Probabilistic graphical models, which combine graph theory with probability theory to represent complex probability distributions, can provide an alternative to logic-based representations [42, 6]. There are primarily two motivations behind probabilistic graphical models. First, even for binary random variables, we need to learn 2^*n*^ − 1 parameters for a distribution of *n* variables. This is unmanageable on many levels: it is computationally difficult to do inference with so many parameters, requires a large amount of memory, and makes it difficult to learn parameters without an unreasonable volume of data [42]. Second, probabilistic graphical models provide important information about independences and the overall structure of the distribution. Probabilistic graphical models were also used as expert systems: *Munin* had a network of more than 1000 nodes to analyze electromyographic data [22], while *PathFinder* assisted medical professionals for the diagnostic of lymph-node pathologies [33] (Figure 1).

**Figure 1:**
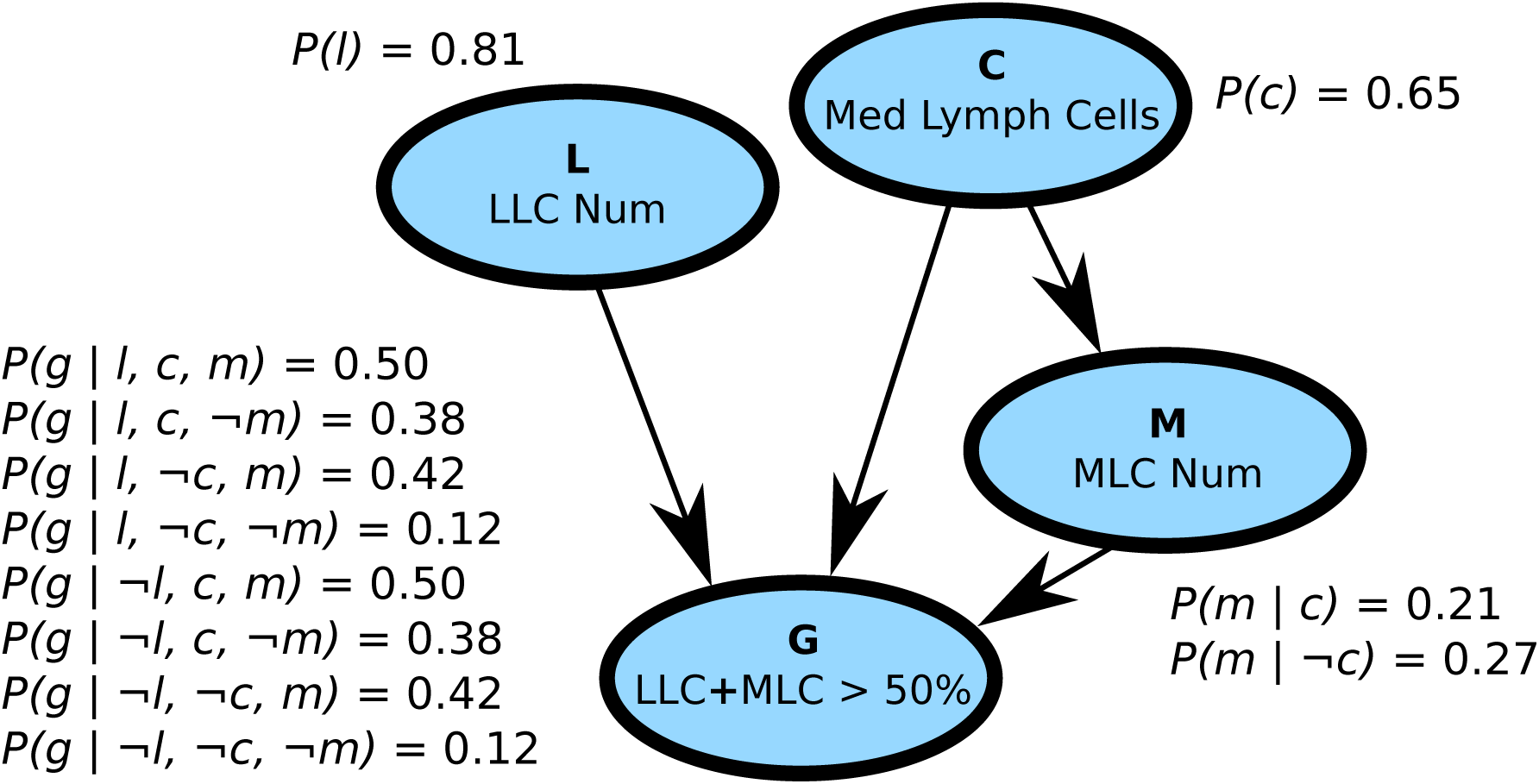
A Bayesian network with four binary variables (the vertices) and possible conditional probability tables. Bayesian networks encode the distribution as directed acyclic graphs such that *P* (**X** = **x**) = Π_*i*_ *P* (*x*_*i*_|*Pa*(*x*_*i*_)), where *Pa*(*x*_*i*_) is the set of parents of variable *x*_*i*_. Because no cycles are allowed, the variables form an ordering so the set *Pa*(*x*_*i*_) can only involve variables already seen on the left of *xi*. Thus, *P* (*a*)*P* (*b*|*a*)*P* (*c*) is a valid Bayesian networks but not *P* (*a*)*P* (*b*|*c*)*P* (*c*|*b*). The four vertices represented here were extracted from *PathFinder*, a Bayesian network with more than 1000 vertices used to help diagnose blood infections [33]. The vertices represent four variables related to blood cells and are denoted by a single character (in bold in the figure): *C, M, L, G*. We denote a positive value with a lowercase letter and a negative value with ¬ (e.g.: *C* = *c, M* = ¬*m*). Since *P* (¬*x*|**y**) = 1 − *P* (*x*|**y**), we need only 2^|*P a*(*x*)|^ parameters per vertex, with |*Pa*(*x*)| being the number of parents of vertex *x*. The structure of Bayesian networks highlights the conditional independence assumptions of the distribution and reduces the number of parameters for learning and inference. As a example query: *P* (*l*, ¬*c, m*, ¬*g*) = *P* (*l*)*P* (¬*c*)*P* (*m*| ¬*c*)*P* (¬*g*|*l*, ¬*c,m*) = 0.81 × (1 − 0.65) × 0.27 × (1 − 0.42) = 0.044. See [17] for a detailed treatment of Bayesian networks and [42] for a more general reference on probabilistic graphical models.

The two key inference problems in probabilistic machine learning are finding the most probable joint state of the unobserved variables (maximum a posteriori, or MAP) and computing conditional probabilities (conditional inference). In a simple presence/absence model for 10 species (*s*_0_, *s*_1_, …, *s*_9_), given that we know the state of species *s*_0_ = *Present, s*_1_ = *Absent, s*_2_ = *Absent*, MAP inference would tell us the most likely state for species *s*_3_, …, *s*_9_, while conditional inference could answer queries such as *P* (*s*_4_ = *Absent*|*s*_0_ = *Present*).

## 2 Markov logic

At this point we have first-order logic, which is capable of manipulating complex logic and mathematical formulas but cannot handle uncertainty, and probabilistic graphical models, which cannot be used to represent mathematical formulas (and thus theories in ecology and evolution) but can handle uncertainty. The limit of first-order logic can be illustrated with our previous example: predators generally have a larger body weight (*Mass*) than their prey, which we expressed in predicate logic as ∀*s*_*x*_, *s*_*y*_ : *PreyOn*(*s*_*x*_, *s*_*y*_) ⇒ *Mass*(*s*_*x*_) > *Mass*(*s*_*y*_), but this is obviously false for some assignments such as *s*_*x*_ : *grey wolf* and *s*_*y*_ : *moose*. However, it is still useful knowledge that underpins many ecological theories [76]. When our domain involves a great number of variables, we should expect useful rules and formulas that are not always true.

A core idea behind many efforts to unify rich logics with probability theory is that formulas can be weighted, with higher values meaning we have greater certainty in the formula. In pure logic, it is impossible to violate a single formula. With weighted formulas, an assignment of concrete values to variables is only *less likely* if it violates formulas, and how much less likely will depend on the weight assigned to the violated formula. The higher the weight of the formula violated, the less likely the assignment is. It is conjectured that all perfect numbers are even (∀*x* : *Perfect*(*x*) ⇒ *Even*(*x*)), thus, if we were to find a single odd perfect number, that formula would be refuted. It makes sense for mathematics but for many disciplines, such as biology, important principles are only expected to be true *most* of the time. If we were to find a single predator smaller than its prey, it would definitely not make our rule useless.

The idea of weighted formulas is not new. Markov logic networks (or just Markov logic), invented a decade ago, allows for logic formulas to be weighted [64, 21]. Similar efforts also use weighted formulas [4, 35]. Markov logic supports algorithms to add weights to existing formulas given a dataset, learn new formulas or revise existing ones, and answer probabilistic queries (MAP or conditional). As a case study, Yoshikawa et al. used Markov logic to understand how events in a document were time-related [78]. Their research is a good case study of interaction between traditional theory-making and artificial intelligence. The formulas they used as a starting point were well-established logic rules to understand temporal expressions. From there, they used Markov logic to weight the rules, adding enough flexibility to their system to beat the best approach of the time. Brouard et al. [13] used Markov logic to understand gene regulatory networks, noting how the resulting model provided clear insights, in contrast to more traditional machine learning techniques. Markov logic greatly simplifies the process of growing a base of knowledge: two research labs with different knowledge bases can simply put all their formulas in a single knowledge base. The only steps required to merge two knowledge bases is to put all the formulas in a single knowledge base and reevaluate the weights.

In a nutshell, a knowledge base in Markov logic ℳ is a set of formulas *f*_0_, *f*_1_, *f*_2_, … along with their weights *w*_0_, *w*_1_, *w*_2_, … :

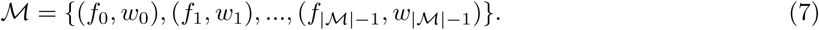

Given constants 𝒞 = {*c*_0_, *c*_1_, *…*, *c*_|*C*|−1_}, ℳ defines a Markov network (an undirected probabilistic graphical model) which can be used to answer probabilistic queries. **Weights** are real numbers in the −∞, ∞ range. The intuition is: the higher the weight associated with a formula, the greater the penalty for violating it (or alternatively: the less likely a possible world is). The **cost** of an assignment is the sum of the weights of the unsatisfied formulas (those that are false). The higher the cost, the less likely the assignment is. Thus, if a variable assignment violates 12 times a formula with a weight of 0.1 and once a formula with a weight of 1.1, while another variable assignment violates a single formula with a weight of 5, the first assignment has a higher likelihood (cost of 2.3 vs 5). Formulas with an infinite weight act like formulas in pure logic: they cannot be violated without setting the probabilities to 0. In short, a knowledge base in pure first-order logic is exactly the same as a knowledge base in Markov logic where all the weights are infinite. In practice, it means mathematical ideas and axioms can easily be added to Markov logic as formulas with an infinite weight. Formulas with weights close to 0 have little effect on the probabilities and the cost of violating them is small. A formula with a negative weight is expected to be false. It is often assumed that all weights are positive real numbers without loss of generality since (*f*, −*w*) ≡ (¬*f, w*). See Jain [38] for a detailed treatment of knowledge engineering with Markov logic. Markov logic can answer queries of complex formulas of the form:

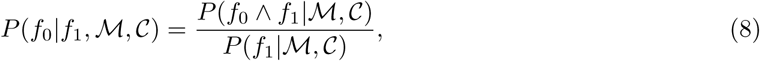

where *f*_0_ and *f*_1_ are first-order logic formulas while ℳ is a weighted knowledge base and 𝒞 a set of constants. It’s important to note that neither *f*_0_ nor *f*_1_ need to be in ℳ. Logical entailment ℳ |= *f* is equivalent to finding *P* (*f*|ℳ) = 1 [21].

We build a small knowledge base for an established ecological theory: the niche model of trophic interactions [76]. The first iteration of the niche model posits that all species are described by a niche position *N* (their body size for instance) in the [0, 1] interval, a diet *D* in the [0, *N*] interval, and a range *R* such that a species preys on all species with a niche in the [*D* − *R/*2, *D* + *R/*2] interval. We can represent these ideas with three formulas:

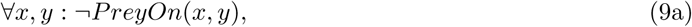

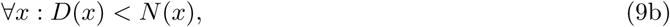

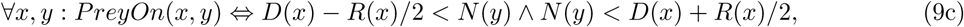

As pure logic, this knowledge base makes little sense. Formula 9a is obviously not true all the time but often is since most pairs of species do not interact. In Markov logic, it is common to have a formula for each lone predicate, painting a rough picture of its marginal probability [21, 38]. We could also add that cannibalism is rare ∀*x* : ¬*PreyOn*(*x, x*) and that predator-prey relationships are generally asymmetrical ∀*x, y* : *PreyOn*(*x, y*) ⇒ ¬*PreyOn*(*y, x*) (although this formula is redundant with the idea that predators are generally larger than their prey). Formulas that are often wrong are assigned a lower weight but can still provide useful information about the system. The second formula says that the diet is smaller than the niche value. The last formula is the niche model: species *x* preys on *y* if and only if species *y*’s niche is within the diet interval of *x*. Using Markov logic and a dataset, we can learn a weight for each formula in the knowledge base. This step alone is useful and provides insights into which formulas hold best in the data. With the resulting weighted knowledge base, we can make probabilistic queries and even attempt to revise the theory automatically. We could find, for example, that the second rule does not apply to parasites or some group and get a revised rule such as ∀*x* : ¬*IsParasite*(*x*) ⇒ *D*(*x*) < *N* (*x*).

## 3 Fuzziness

First-order logic provides a formal language for expressing mathematical and logical ideas while probability theory provides a framework for reasoning about uncertainty. A third dimension often found in discussions on unifying logic with probability is fuzziness. A struggle with applying logic to ecology is that all predicates are either true or false. Even probabilistic logics like Markov logic define a distribution over *binary* predicates. Going back to Rosenzweig-MacArthur (eq. 3), this formula’s weight in Markov logic is almost certainly going to be zero since it’s never *exactly* right. If the Rosenzweig-MacArthur equation predicts a population size of 94 and we observe 93, the formula is false. Weighted formulas help us understand *how often a formula is true*, but in the end the formula has to give a binary truth value: true or false, there is no place for nuance. Logicians studied more flexible logics where truth is a real number in the [0, 1] range. These logics are said to be “infinitely many-valued” or “fuzzy”. In this setting: 0 is false, 1 is true, and everything in-between is used to denote nuances of truth [79, 7]. Predicates returning truth values in the [0, 1] range are called **fuzzy predicates**, while standard predicates returning *false, true* are said to be bivalent. To show fuzziness in action, let’s look at a simple formula that says that small populations experience exponential growth:

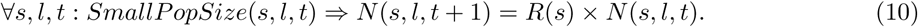

Variables *s, l, t* respectively denote a species, a location, and time. Function *N* returns the population size of a species at a specific location and time while function *R* returns the growth rate of the species. The predicates *SmallPopSize* and = are both problematic from a bivalent perspective. Equality poses problem for the same reason it did with the Rosenzweig-MacArthur example: we do not expect this formula to be exactly right. The notion of a small population size should also be flexible, yet logic forces us to determine a strict threshold where *SmallPopSize* will change from true to false. Using truth values in the [0, 1] range makes it possible to have a wide range of nuances for both *SmallPopSize* and equality. See table 2 for the definitions of fuzzy logic connectives.

**Table 2:**
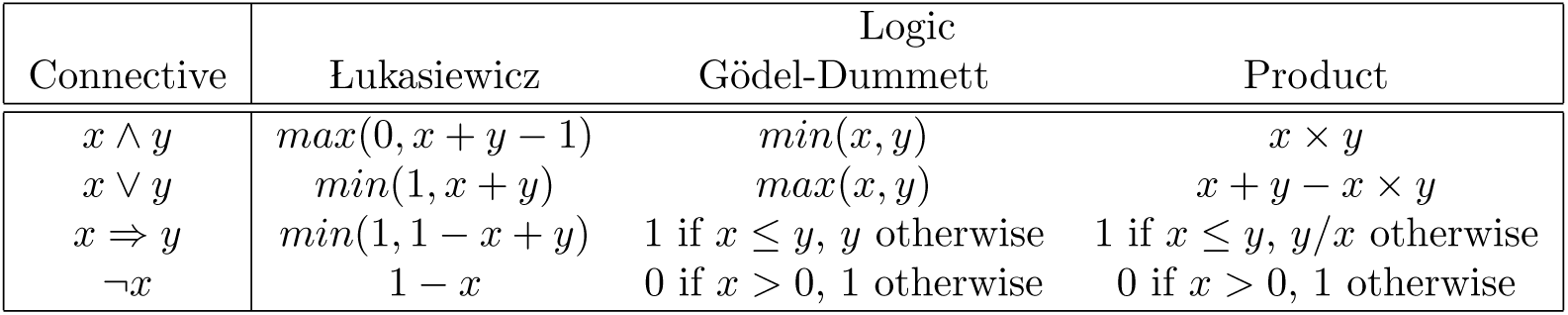
Definitions of logic connectives for the three main fuzzy logics. These three logics are said to be normal, meaning they behave exactly like classical logic when restricted to truth values of 0 (false) and 1 (true). When truth values are between 0 and 1, these logics will often behave differently than classical logic. For example, in both classical and L ukasiewicz logics, ¬¬*x* ≡ *x*, but it is not the case for Gödel-Dummett and Product logics (unless *x* ∈ {0, 1}). Another example is that conjunction and disjunction are idempotent in classical and Gödel-Dummett logics, meaning *x* ∧ *x* ≡ *x* and *x* ∨ *x* ≡ *x*, but it is not the case for Łukasiewicz and Product logics. See [7] for a detailed explanation of how the connectives are defined.

Fuzzy logic is not a replacement for probability theory. The most interesting aspect of fuzzy logic is how it interacts with probability theory to form truly flexible languages. For examples, fuzzy predicates are used in both probabilistic soft logic [40, 4] and deep learning approaches to predicate logic [80, 35]. Hybrid Markov logic [74, 21] extends Markov logic by allowing not only weighted formulas but terms like soft equality, which applies a Gaussian penalty to deviations from equality. While not exactly a full integration of fuzzy logic into Markov logic, soft equality behaves in a similar matter and is a good fit for formulas like the Rosenzweig-MacArthur system or our previous example with exponential growth. Hybrid Markov logic is not as well-developed as standard Markov logic, for example there are no algorithms to learn new formulas from data. On the other hand, Hybrid Markov logic solves many of the problems caused by bivalent predicates while retaining the ability to answer conditional queries. In the next section we’ll explore hybrid Markov logic and its application to an ecological dataset. Several languages for reasoning have combined fuzziness with probability or logic (Figure 2). It has been argued that, in the context of Bayesian reasoning, fuzziness plays an important role in bridging logic with probability [55, 37]. However, how to effectively combine rich logics with probability theory remains an open question, as is the role of fuzziness.

**Figure 2:**
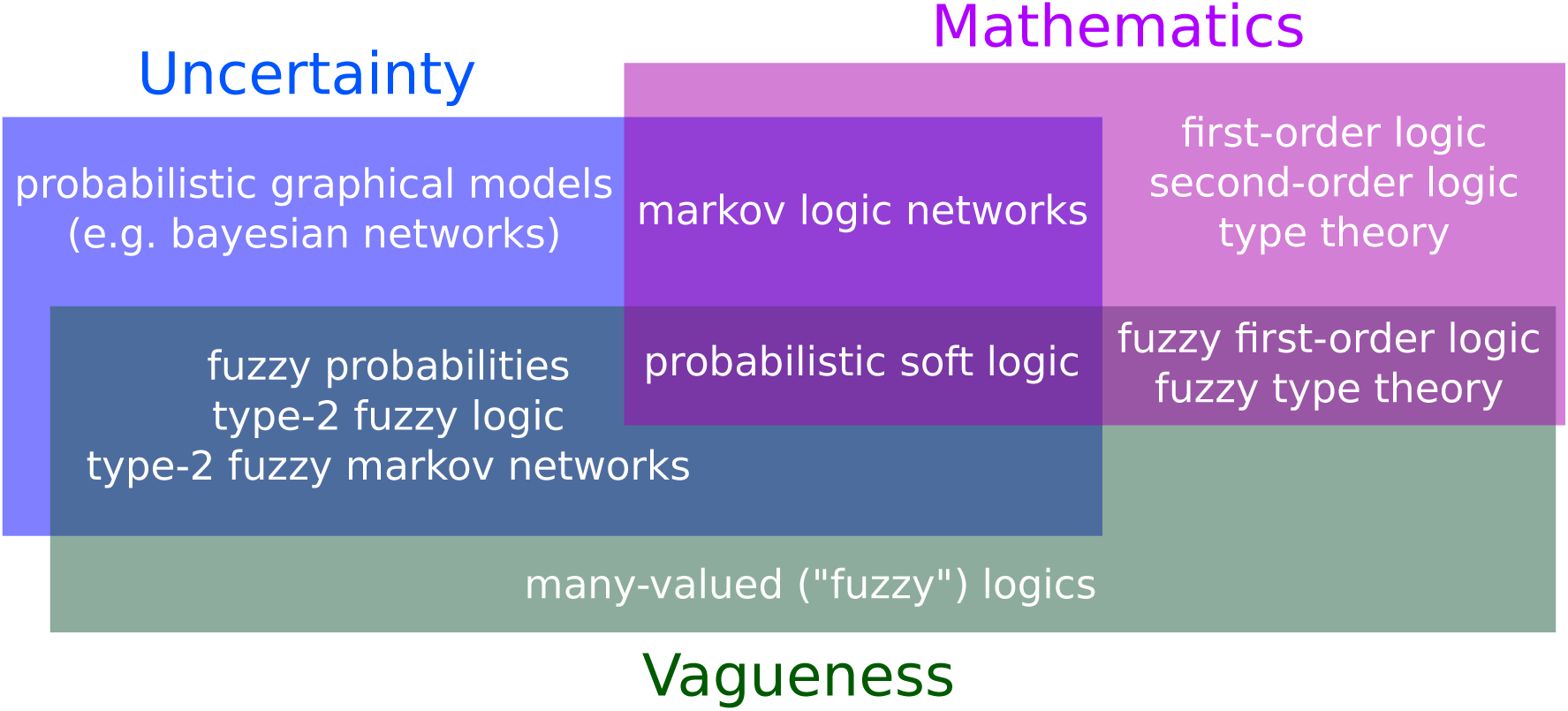
Various languages and their ability to model uncertainty, vagueness, and mathematics (the size of the rectangles has no meaning). **In the blue rectangle**: languages capable of handling uncertainty. Probabilistic graphical models combine probability theory with graph theory to represent complex distributions [42]. Alternatives to probability theory for reasoning about uncertainty include possibility theory and Dempster-Shafer belief functions, see [30] for an extended discussion. **In the green rectangle**: Fuzzy logic extends standard logic by allowing truth values to be anywhere in the [0, 1] interval. Fuzziness models vagueness and is particularly popular in linguistics, engineering, and bioinformatics, where complex concepts and measures tend to be vague by nature. See [44] for a detailed comparison of probability and fuzziness. **In the purple rectangle**: languages capable of modelling mathematical formulas. It is important to note that while first-order logic is expressive enough to express a large class of mathematical ideas, many languages rely on a restricted from of first-order logic without functions. Alone, these languages are not powerful enough to express scientific ideas, we must thus focus on what lies at their intersection. Type-2 Fuzzy Logic is a fast-expanding [67, 50] extension to fuzzy logic, which, in a nutshell, models uncertainty by considering the truth value itself to be fuzzy [51, 81]. Markov logic networks [64, 21] extends predicate logic with weights to unify probability theory with logic. Probabilistic soft logic [40, 3] also has formulas with weights, but allows the predicates to be fuzzy, i.e. have truth values in the [0, 1] interval. Some recent deep learning studies also combine all three aspects [25, 35].

## 4 Markov logic and the Salix tritrophic system

The primary goal of unifying logic and probability is to be able to grow knowledge bases of formulas in a clear, precise language. For Markov logic, it means a set of formulas in first-order logic. For this example, we used Markov logic to build a knowledge base for ecological interactions around the Salix dataset [43]. The Salix dataset has 126 parasites, 96 species of gallers (insects), and 52 species of salix, forming a tritrophic ecological network (*Parasite* → *Galler* → *Salix*). Furthermore, we have partial phylogenetic information for the species, their presence/absence in 374 locations, interactions, and some environmental information on the locations. To fully illustrate the strengths and limits of Markov logic in this setting, we will not limit ourselves to the data available for this particular dataset (e.g. we do not have body mass for all species).

Data in first-order logic can be organized as a set of tables (one for each predicate). For our example, we have a table named *PreyOnAt* with three columns (its arguments) and a table named *IsParasitoid* with only one column. This format implies the closed-world assumption: if an entry is not found, it is false (see table 4 for an example). For this problem we defined several functions and predicates to describe everything from predator-prey relationships, whether pairs of species often co-occurred, along with information on locations such as humidity, precipitation, and temperature (see table 3). We ran the basic learning algorithm from Alchemy-2 [64], which is used both to learn new formulas and weight them. The weights are listed at the end of each formula. We use the “?” character at the end of the formula involving data that were unavailable for this dataset (and thus, we could not learn the weight). Here’s a sample of a knowledge base where the first three formulas were learned directly from our dataset and the last two serve as example for Hybrid Markov logic:

**Table 3:** Predicates and functions used for the Salix example. A predicate is simply a function mapping to a boolean value (false or true, denoted *bool*). ℕ stands for natural numbers (0, 1, 2, …) while ℝ stands for real numbers, and [0, 1] is a shorthand for a real number in the [0, 1] range. We must often force continuous values into boolean values. For example, *HighTemperature* and *CloselyRelated* both require arbitrary cutoffs, often the line between *true* and *false* is set at the mean. Recent languages push for greater integration with fuzziness, which would allow predicates to take any values in the [0, 1] range.

| Functions | Meaning |
| --- | --- |
| $PPreyOn : species \times species \mapsto [0, 1]$ | Probability that a species preys on another |
| $PreyOn : species \times species \mapsto bool$ | Predator-prey relationship |
| $PreyOnAt : species \times species \times location \mapsto bool$ | Predator-prey relationship at a given location |
| $PresenceAt : species \times location \mapsto bool$ | Presence of a species at a location |
| $IsParasite : species \mapsto bool$ | Whether the species is a parasite |
| $IsGaller : species \mapsto bool$ | Whether the species is a galler |
| $IsSalix : species \mapsto bool$ | Whether the species is a salix |
| $CloselyRelated : species \times species \mapsto bool$ | Whether two species are closely related |
| $Occ : species \mapsto \{location\}$ | Set of locations where a species is found |
| $Cooccurrence : species \times species \mapsto \mathbb{R}^+$ | Proportion of locations where the species co-occur |
| $HighCooccurrence : species \times species \mapsto bool$ | Pair of species with high co-occurrence |
| $HighTemperature : location \mapsto bool$ | Location with above-average temperature |
| $T : \{location\} \mapsto \mathbb{R}$ | Mean temperature for a set of locations |
| $Mass : species \mapsto \mathbb{R}^+$ | Mean adult body mass for a species |
| $FoodWeb : location \mapsto Graph$ | Food web at a given location |
| $Connectance : Graph \mapsto \mathbb{R}^+$ | $Edges/Vertices^2$ |
| $SpeciesRichness : Graph \mapsto \mathbb{N}$ | Number of species in the food web |
| $N : species \mapsto \mathbb{R}^+$ | Niche of species per [75] |
| $C : species \mapsto \mathbb{R}^+$ | Diet of the species per [75] |
| $R : species \mapsto \mathbb{R}^+$ | Range of species' diet per [75] |

**Table 4:** A sample of three tables for the Salix dataset [43]. Species are denoted by the first six letters of their names while sites are numbered from 1 to 374. Data in first-order logic is often organized in tables with one table per predicate and where entries represent true values while absent combinations are assumed to be false. For example, given this sample, *HighTemp*(*Site*006) is true while *HighTemp*(*Site*001) would be false. The full data formatted for Alchemy-2 [64] is provided as supplementary material.

| *PreyOnAt* |  |  | *IsParasitoid* | *HighTemperature* |
| --- | --- | --- | --- | --- |
| Amorri | Ovesic | Site060 | Ppecti | Site006 |
| Chalci | Halien | Site116 | Psoemi | Site311 |
| Ireuni | Hpolit | Site291 | Tspone | Site296 |
| Eacicu | Ovimin | Site121 | Tsptwo | Site183 |
| ... | ... | ... | ... | ... |

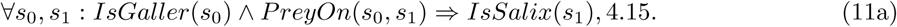

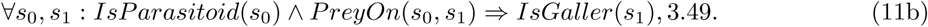

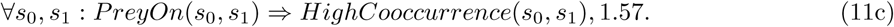

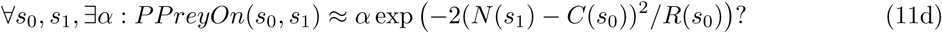

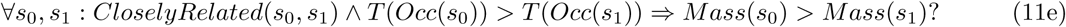

The first two formulas correctly define the tritrophic relationship between parasites, galler and salix, while the third shows a solid, but not as strong, relationship between predation and co-occurence. Formula 11d would require hybrid Markov logic and a fuzzy predicate ≈.

Integration of macroecology and food web ecology may rely on a better understanding of macroecological rules [5]. These rules are easy to express with first-order logic, for example equation 11e is a formulation of Bergmann’s rule. We also used the learning algorithm to test whether closely related species had similar prey, but the weight attributed to the formula was almost zero, telling us the formula was right as often as it was wrong:

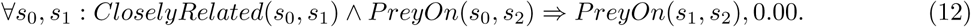

This example shows both the promise and the current issues with hybrid logic-probabilistic techniques. Many of the predicates would benefit from being fuzzy, for example, *PreyOn* should take different values depending on how often predation occurs. We also had to use arbitrary cut-offs for predicates like *CloselyRelated* and *HighTemperature*. Fortunately, many recent approaches integrate logic with both fuzziness and probability theory [1, 35, 4]. Weights are useful to understand which relationship is strong in the data, and this example shows the beginning of a knowledge base for food web ecology. The next step would be to discover new formulas, whether manually or using machine learning algorithms, and add data to revise the weights. If a formula involves a predicate operating on food webs and we want to apply our knowledge base to a dataset without food webs, this formula will simply be ignored (because it won’t have grounded predicates to evaluate it; see section 1). This is a strong advantage of this knowledge representation: our little knowledge base here can be used as a basis for any other ecological datasets even if they quite different. With time, it’s possible to grow an increasingly connected knowledge base, linking various ideas from different fields together.

## 5 Bayesian Higher-Order Probabilistic Programming

Artificial Intelligence has a long history with first-order logic [66] but type theory (or higher-order logic), a more expressive logic, is currently more popular both as a tool to formalize mathematics and as foundation for programming languages. We explored hybrid approaches based on first-order logic and, for this section, we’ll briefly discuss Bayesian Higher-Order Probabilistic Programming (BHOPP) along with its relationship with type theory. Probabilistic programming languages are programming languages built to describe probabilistic models and simplify the inference process. Stan [15] and BUGS [47] are two popular examples of probabilistic programming languages used for Bayesian inference, but even more flexible languages for Bayesian probabilistic programming have recently emerged. These languages, like Church [29] and Anglican [77], accept higher-order constructs (that is: functions accepting other functions as arguments). The ambition is that “ultimately we would like simply to be able to do probabilistic programming using any existing programming language as the modeling language” [73].

First-order logic allowed us to model intricate theories but, in practice, almost all modern systems used to formalize mathematics are based on type theory (higher-order logic) [56]. The “first” in first-order logic refers to the limitation that quantification can only be done on individual elements of a set, but not on higher-order structures like sets, predicates, or functions. As a consequence, several important concepts in mathematics cannot be formalized directly with first-order logic. Since type theory supports higher-order quantification, it is used as a foundation to reason about mathematics. Coq, HOL, HOL Light, and Microsoft’s LEAN are all popular languages for automated theorem proving based on different forms of type theory [48, 32, 18]. Programming languages in general, not just those targeted at mathematicians, tend to also rely on type theory as foundation [58]. See [24, 56] for an introduction to type theory. Here is where it gets confusing: the *higher* in higher-order logic has a different meaning than in *higher-order probabilistic programming* and yet, Bayesian higher-order probabilistic programming languages (BHOPPL) may hold the key to sound inference mixed with type theory. In BHOPPL, *higher-order* means functions can take functions as arguments, a common capability of modern programming languages. This is necessary for higher-order logic but not sufficient. Where it gets exciting is that a lot of progress is being made in framing BHOPPL in the language of type theory [12]. In effect, it would bring Bayesian and higher-order logic reasoning together.

Furthermore, software-wise, BHOPPLs are well ahead of the approaches described in previous sections such as hybrid Markov logic networks. Current higher-order probabilistic programming languages operate on variants of well-known languages: Anglican is based on Clojure [77], Pyro is based on Python [9], Turing.jl uses Julia [26]. Many BHOPPLs have been designed to exploit the high-performance architecture developed for deep learning such as distributed systems of GPUs (graphics cards). GPUs have been important in the development of fast learning and inference in deep learning [28]. Pyro [9] is a BHOPPL built on top of PyTorch, one of the most popular frameworks for deep learning, allowing computation to be distributed on systems of GPUs. In contrast, there are no open-source implementations of Markov logic networks running on GPUs. The main downside of BHOPPLs is that, while in theory they may support the richer logics used to formalize modern mathematics, in practice higher-order probability theory is itself not well understood. This is an active research topic [73] but formalization faces serious issues. For one, there are incompatibilities with the standard measure-theoretic foundation of probability theory, which may require rethinking how probability theory is formulated [11, 70, 69, 34, 68]. First-order logic is among the most studied formal languages, making it easy to use a first-order knowledge base with various software. The current informal nature of BHOPPLs make them hard to recommend for the synthesis of knowledge in ecology and evolution, even though they may very well hold the the most potential.

## 6 Where’s our unreasonably effective paradigm?

Legitimate abstractions can often obfuscate how much various subfields are related. Natural selection is a good example. Many formulas in population genetics rely on fitness. Nobody disputes the usefulness of this abstraction, it allows us to think about changes in populations without worrying whether selection is caused by predation or climate change. On the other hand, fitness has also allowed the development of theoretical population genetics to evolve almost independently of ecology. There is a realization that much of the complexity of evolution is related to how selection varies in time and space, which puts evolution in ecology’s backyard [8]. Achieving Lewis’ goal of formalization would not prevent the use of fitness, but having formulas with fitness cohabiting with formulas explaining the components of fitness would implicitly link ecology and evolution. This goes in both directions: what are the consequences of new discoveries on speciation and adaptive radiations on the formation of metacommunities? How can community dynamics explain the extinction and persistence of new species? If there isn’t a single theory of biodiversity, the imperative is to understand biodiversity as a system of theories. Given the scope of ecology and evolution and the vast number of theories involved, it seems difficult to achieve a holistic understanding without some sort of formal system to see how the pieces of the puzzle fit together. Connolly et al. noted how theories for metacommunities were divided between those derived from first principles and those based on statistical methods [16]. In systems unifying rich logics with a probabilistic representation, this distinction does not exist, theories are fully realized as symbolic and statistical entities. Efforts to bring theories in ecology and evolution into a formal setting should be primarily seen as an attempt to put them in context, to force us to be explicit about our assumptions and see how our ideas interact [71].

Despite recent progress at the frontier of logic and probability, there are still practical and theoretical issues to overcome to make a large database of knowledge for ecology and evolution possible. Inference can be difficult in rich knowledge representations, not all methods have robust open-source implementations, and some approaches such as Bayesian higher-order probabilistic programming are themselves not well understood. Plus, while mathematicians benefit from decades of experience making large databases of theorems, there have been no such efforts for ecology and evolution. Lewis’ case for the formalization is worth repeating: “when theories are partially formalized […] the intra- and interworkings of theories become more clearly visible, and the total structure of the discipline becomes more evident” [45]. This vision might soon become reality thanks to increased access to data in evolution and evolution and recent advances at the frontier of logic and probability. Given the pressing need to understand a declining biodiversity, ecologists and evolutionary biologists should be at the forefront of the efforts to organize theories in unified knowledge bases.

## Supporting information

Data for Salix case study

## 7 Acknowledgements

We thank three anonymous reviewers for their helpful comments. PDP has been funded by an Alexander Graham Bell Graduate Scholarship from the National Sciences and Engineering Research Council of Canada, an Azure for Research award from Microsoft, and benefited from the Hardware Donation Program from NVIDIA. DG is funded by the Canada Research Chair program and NSERC Discovery grant. TP is funded by an NSERC Discovery grant and an FQRNT Nouveau Chercheur grant.

